# Multi-agent AI System for High Quality Metadata Curation at Scale

**DOI:** 10.1101/2025.06.10.658658

**Authors:** Rajdeep Mondal, Manimala Sen, Sahana Sengupta, Writtik Maity, Shyam Palapetta, Nobal Kishor Dhruw, Abhishek Jha

## Abstract

High-quality metadata is essential for downstream AI applications, yet metadata curation remains a persistent bottleneck in biomedical research. The core challenge lies in balancing quality and scalability. Manual curation delivers high quality, reliable annotations but is time-intensive and non-scalable; automated approaches, including those based on natural language processing, can scale but often fall short in accuracy and completeness. This tradeoff is particularly severe in public datasets, where metadata is frequently distributed across multiple sources for a single dataset. A notable example includes multi-omics datasets available on GEO and similar public repositories, which are often accompanied by associated publications. These supplementary sources can be leveraged to substantially enhance metadata quality, thereby supporting downstream applications such as classifier model development or the identification of biologically relevant cohorts, etc.

We present a first-in-class multi-agent AI system that bridges the gap, achieving both high quality and scalability in metadata curation. Built on large-language models (LLMs), our system orchestrates a set of specialized agents that collaboratively extract, normalize, and infer critical metadata fields such as tissue, disease, cell line, sampling site, demographics, and experimental context from GEO entries, associated publications. A central orchestrator agent delegates tasks such as data retrieval, document parsing, ontology mapping, and context inference to expert sub-agents, enabling scalable, end-to-end automation.

Applied to GEO data sets, a notoriously difficult metadata domain, our system achieves a 93% recall on average across 23 key fields (all original terms, normalized terms, and corresponding ontology ids) that include information about disease, tissue, treatment, donor-related information, outperforming existing automated baselines and approaches expert level quality. We also present a system that can easily scale to curate hundreds of metadata fields of interest with similar precision. This work demonstrates that an LLM-based multi-agent architecture can overcome traditional trade-offs in metadata curation, enabling both precision and scale, and offers a promising path forward for curating large public biomedical repositories for downstream AI applications.

## 1. Introduction

Artificial Intelligence has increasingly become an integral part of modern drug discovery, enabling the identification of hidden patterns and mechanistic insights that traditional methods often miss. Using machine learning and deep learning, AI integrates diverse biomedical data from omics to clinical records to facilitate a deeper understanding of disease biology and therapeutic response. Its applications span biomarker and target discovery, indication prioritization, de novo molecule generation, pharmacokinetic modeling, drug– target interaction prediction, and clinical trial optimization. Together, these advancements are expediting the drug development pipeline and contributing to the growing number of AI-derived candidates progressing into clinical trials [1].

A cornerstone of effective data utilization for AI applications is the availability of high-quality metadata, which provides essential descriptive information on samples, experimental design, data processing procedure, etc. Comprehensive and accurate metadata are paramount for ensuring the findability, accessibility, interoperability, and reusability (FAIR) of biological datasets, allowing researchers to contextualize the data, evaluate its quality, and integrate it with other datasets for comprehensive analysis [2]. In fact, the potential for data reuse and the efficiency of scientific discovery are directly linked to the quality and completeness of associated metadata. In contrast, inadequate metadata can significantly hinder the extraction of valuable knowledge from these datasets, acting as a major bottleneck in the research process [3].

However, manual metadata curation for the rapidly growing volume of datasets, particularly within public repositories such as the Gene Expression Omnibus (GEO), poses substantial challenges due to inconsistencies, inaccuracies, and non-compliance with FAIR quality standards. This process of making the data useful is inherently labor intensive and requires expert domain knowledge and a deep understanding of the scientific context. GEO, while a vital resource, exhibits considerable data heterogeneity, inconsistencies in experimental protocols and sample handling, and discrepancies in data submission practices by authors. Furthermore, critical metadata fields are frequently absent from GEO records, necessitating their extraction from associated publications and supplementary materials, a task that is both time-consuming and difficult to automate effectively. Scientific publications can be extensive, and supplementary files are often provided in a variety of formats, adding to the complexity of automated processing. The limitations of manual curation in the face of increasing data volumes and the inherent inconsistencies within public repositories highlight the urgent need for automated solutions capable of efficient and accurate metadata extraction [4].

Several strategies have been proposed to enhance the curation of metadata linked to publicly available studies in the GEO repository. These approaches generally fall into three main categories: (1) manual curation, (2) automated methods using natural language processing (NLP), and (3) metadata inference based on gene expression data itself [5]. Recently, multi-agent systems like BioAgents have been built to assist users in the design, development, and troubleshooting of complex bioinformatic pipelines [6].

Multi-agent AI systems offer a promising paradigm for addressing this complex problem by harnessing the collective capabilities of multiple specialized agents working in tandem. This paper introduces a first-in-class multi-agent AI system designed for the automated curation of metadata fields for single-cell RNA sequencing datasets obtained from public sources like GEO, as well as from associated publications and supplementary materials. The system incorporates specialized agents designed for various tasks, including data acquisition, scientific papers processing, managing disparate supplementary file formats, extracting relevant entities, and performing reasoning. It employs specific strategies to navigate challenges such as the processing of lengthy documents, the management of varied file types, and the resolution of inconsistent annotations. The primary objectives of this paper are to detail the design and implementation of this multiagent AI system, to discuss the specific challenges in metadata curation, and to outline its potential benefits for the broader biomedical research community.

## 2. Related Work

The development of automated systems for the curation of biomedical metadata is based on decades of research in computational biology, natural language processing, and knowledge representation [2]. This section synthesizes critical advancements across key interrelated domains that collectively enable multi-agent approaches to metadata harmonization and ontology alignment.

### 2.1 Metadata Heterogeneity in Genomic Repositories

Modern genomic repositories face fundamental challenges in metadata consistency due to heterogeneous reporting practices between research groups [7]. Early standardization efforts such as MetaSRA demonstrated the feasibility of ontology mapping through computational pipelines, successfully mapping samples to terms in biomedical ontologies including Uberon, Disease Ontology, and Cell Ontology [8]. MetaSRA automated the normalization of human sample-specific metadata from the Sequence Read Archive, following a schema inspired by the ENCODE project [9].

Subsequent developments have focused on machine learning approaches to address metadata inconsistencies [10]. Recent work has explored AI-driven curation systems, exemplified by SRAgent from scBaseCamp, which employs hierarchical agent workflows to systematically identify and integrate single cell RNA sequencing datasets from public repositories [11].

### 2.2. Large Language Models in Biomedical NLP

The advent of transformer-based language models has transformed biomedical information extraction capabilities [12]. Domain-specific adaptations such as BioBERT and PubMedBERT demonstrated that masked language model pretraining on biomedical corpora achieves improved performance on named entity recognition tasks, with BioBERT showing 0. 62% F1 score improvement for biomedical NER and 2.80% improvement for relation extraction compared to general BERT models [12].

Generative approaches have emerged through decoder-based architectures such as BioGPT, which employs a GPT-2 architecture with 347M parameters trained on 15M PubMed abstracts [13]. Recent applications demonstrate the potential of fine-tuned large-language models for specialized biomedical tasks, with ChIP-GPT achieving a precision of 90 - 94 % in extracting metadata from chromatin immunoprecipitation experiments when trained with 100 examples [14].

Furthermore, critical to modern approaches is the integration of retrieval-augmented generation (RAG) techniques that ground language model outputs in verified knowledge bases [15].

### 2.3. Multi-Agent Systems for Complex Workflows

The computational complexity of metadata curation has motivated architectures that surpass traditional monolithic pipelines [6]. Multi-agent systems have emerged as a paradigm for orchestrating distributed intelligence across specialized components, particularly in domains requiring heterogeneous data integration. BioAgents established a framework that combines retrieval-augmented language models with domain-specific reasoning modules, demonstrating performance comparable to that of human experts in conceptual genomics tasks [6].

### 2.4. Reasoning Frameworks for Ambiguity Resolution

Effective metadata curation requires sophisticated reasoning capabilities to resolve inherent ambiguities in biomedical reporting [16]. Chain-of-thought prompting has proven effective for multistep inference tasks, with some biomedical adaptations achieving accuracy levels around 81% for diagnostic reasoning tasks [16]. The tree-of-thought framework improves this by maintaining multiple concurrent reasoning paths, offering potential improvements for complex problem solving scenarios [17].

### 2.5. Ontology Mapping and Semantic Integration

The mapping of biomedical ontologies remains fundamental for the interoperability of metadata [18]. The BioHAN framework demonstrated that hyperbolic graph attention networks can effectively perform cross-ontology mapping by jointly modeling terminological and structural semantics [18]. Complementing this, SapBERT introduced self-supervised pretraining in UMLS concept pairs to learn dense biomedical entity representations, achieving state-of-the-art performance on medical entity linking benchmarks [19].

## 3. Methods

### 3.1. Curation Objective and Schema Design

The objective of this work is to develop a highly accurate and scalable system for automated metadata curation from public transcriptomics datasets, with a focus on datasets archived in the Gene Expression Omnibus (GEO), SCP, ArrayExpress, etc. Manual curation has long been the gold standard for metadata accuracy, but its limitations in throughput and consistency render it insufficient for the scale and diversity of modern omics datasets. Our approach aims to combine the contextual understanding and reasoning capabilities of large language models (LLMs) with a multi-agent design that enables task specialization, auditability, and modular reuse.

To standardize curation across diverse studies, we adopted a modified version of the schema defined by the Chan Zuckerberg Initiative’s CELLxGENE Discover platform (version 5.2.0) as the foundation [20]. This schema provides a structured vocabulary for describing key biological and experimental attributes associated with single-cell RNA sequencing (scRNA-seq) datasets. However, the original schema does not fully account for certain high-value metadata fields such as cell line, treatment, and drug exposure that are frequently reported in publications but inconsistently reflect in public repositories like GEO. To address this gap, we extended the schema to incorporate these fields along with ontology-linked identifiers, thereby enabling more complete downstream data integration and reuse.

The resulting schema comprises both the study-level (GSE) and sample-level (GSM) metadata attributes. At the sample level, nine fields were selected for systematic evaluation based on their relevance to biological interpretation and the availability of the corresponding ontology standards. These include organism, tissue, cell line, disease, developmental stage, sex, self-reported ethnicity, tissue type, and donor ID. For each field, the system is designed to output (i) the raw value as reported in the source (e.g., GEO page, supplementary file, or publication text), (ii) an ontology normalized term, and (iii) a stable ontology identifier (e.g., MONDO for diseases, UBERON for anatomical terms, PATO for phenotypic sex, and HANCESTRO for ancestry terms). The study level fields consist of the GSM ids, DOIs, PMIDs, etc. This schema shown in **Supplementary Table 1** not only provides a common format for representing sample annotations, but also facilitates semantic interoperability across datasets. Inclusion of ontology term identifiers allows harmonized querying and supports downstream applications such as cross-study meta-analysis, patient stratification, and training of machine learning models. Throughout the pipeline, our system maintains both the original author-provided metadata and its normalized counterpart, enabling reversible mappings and flexible integration with external systems that may rely on alternative ontologies.

**Table 1.** Reviewer Concordance for Curated Metadata Labels. This table summarizes the reviewer concordance for a subset of key metadata fields manually evaluated during system validation.

| Label Name | Concordance |
| --- | --- |
| Disease | 1.000 |
| Tissue | 0.934 |
| Tissue Type | 1.000 |
| Gender | 0.970 |

To support future extensibility, the schema design includes optional fields and structured placeholders for recording provenance information, such as the exact sentence, table cell, or GEO field from which a given annotation was derived. This dual-layered representation, linking raw evidence to standardized knowledge, is central to the transparency and trustworthiness of our approach and lays the foundation for active learning and human-in-the-loop refinement in future deployments.

### 3.2. Dataset Collection and Preprocessing

#### 3.2.1. Selection of GEO studies

To evaluate the performance and scalability of our multiagent curation system, we curated a benchmark corpus of single-cell transcriptomic studies from the Gene Expression Omnibus (GEO). Studies were selected based on the following criteria: (i) the study organism must be *Homo sapiens*, (ii) the sequencing technology must be a single-cell or singlenucleus RNA-Seq, and (iii) a corresponding full-text publication must be accessible through PubMed Central or publisher-provided links and that the curated data was also present in CELLxGENE. Studies were further prioritized to ensure diversity in biological context, assay technology (e.g., 10x Genomics, SMART-Seq) and metadata completeness. Our final corpus consisted of 78 entries in the GEO series (GSE) that encompasses 4652 samples (GSMs) corresponding to 42 distinct peer-reviewed publications (**Supplementary Table 2**). In total, this data set included more than 120 million metadata tokens in primary GEO records, full-text articles, and supplementary files, offering a realistic and challenging testbed for metadata extraction. The selected corpus includes studies drawn from consortium-scale initiatives as well as smaller investigator-led datasets to reflect the range of reporting practices in the field.

#### 3.2.2. Retrieval of Source Documents

For each selected GSE, we retrieve study and sample-level metadata using NCBI E-utilities [21]. These records provided structured fields such as title, summary, overall design, and sample-specific characteristics strings. We recorded and parsed all publication identifiers (PMIDs, PMCIDs, DOIs) embedded in the metadata to link each dataset to its associated article.

Full-text publications were retrieved using a hierarchical access strategy. When a PubMed Central ID (PMCID) was available, the full text was downloaded through the Entrez EFetch utility [21]. Otherwise, we attempted to resolve the DOI through cross-ref or direct publisher APIs. For articles behind paywalls, we defaulted to using the abstract and any GEO-provided summaries, although such cases were rare in our corpus. Each retrieved article was converted to plain text and segmented into canonical sections (e.g., Introduction, Methods, Results, Discussion) to enable targeted querying by downstream agents.

In parallel, all supplementary files referenced in the publication were downloaded programmatically. These included structured formats such as Excel (XLS/XLSX), comma- or tab-separated files (CSV/TSV/TXT) and unstructured formats such as PDF and DOCX.

#### 3.2.3. Text and Table Processing

All retrieved publications were first parsed into their canonical sections Abstract, Introduction, Materials and Methods, Results, Discussion, and References - and stored in a structured JSON format. Each section was treated as an independent unit of information; no further categorization was applied beyond this level. When an agent required information from a publication, it selectively queried only the relevant section, thereby preserving both context specificity and efficiency. This design choice also allowed the system to respect the natural boundaries of scientific discourse as intended by the authors while reducing noise from unrelated sections.

For the extraction of structured data from supplementary materials, we implemented a specialized pipeline tailored for high-accuracy table recognition and parsing from PDF documents. The process began with a deterministic program that iterated through every page of each supplementary PDF and identified candidate table pages. This identification step was rule-based and did not involve any large language models (LLMs); instead, it relied on visual and layout cues such as gridlines, consistent row spacing, and table-like text blocks to isolate pages that contained tabular structures.

Pages flagged as containing tables were then saved as high-resolution images and passed to a visual LLM with multimodal reasoning capabilities in this case, GPT-4.1 with image input support. The model was prompted to extract the contents of the table and return a structured JSON object. This object included both the table data and associated metadata, such as the table title (if available), the page number of the original document, and any relevant units or notes embedded in the caption or header rows.

In parallel, we develop a tagging mechanism to label tables that contain demographic information about study subjects or donors. These tables, often buried in supplementary files, are critical for curating fields such as sex, age, ethnicity, donor ID, and developmental stage. The tables identified by the visual model as referring to participants or donor cohorts were explicitly tagged with a donor metadata = true flag, allowing downstream agents responsible for demographic field extraction to prioritize these high-signal sources.

The output of this pipeline was a set of machine-readable JSON files for each parsed table, linked back to their source page and file, and annotated for downstream utility. Together with segmented publication sections and structured GEO metadata, these JSON encoded tables were integrated into each sample’s raw evidence bundle forming a comprehensive input layer for the multi-agent reasoning system.

### 3.3. System Architecture: Multi-Agent Curation Framework

#### 3.3.1. Overview of Agentic Design

To enable accurate and scalable curation of datasets from Gene Expression Omnibus (GEO), we developed a modular, multi-agent system built around cognitive reasoning and structured validation. The architecture is centered on a six-stage processing pipeline that integrates both chain-of-thought [16] and tree-of-thought [17] reasoning paradigms. These allow the system to explore multiple interpretation paths simultaneously, compare outcomes, and converge on high-confidence results through iterative refinement and consensus.

The system is designed for composability and auditability: deterministic preprocessing modules handle data ingestion and formatting, while dedicated reasoning agents - instantiated using GPT-4.1 with function-calling takes over complex extraction, mapping, and normalization tasks. This separation of concerns enables scalable parallel execution, finegrained quality control, and transparent decision-making at each step.

Ontology normalization is handled by a dedicated subsystem that combines dense semantic retrieval (through finetuned SapBERT and PubMedBERT embeddings), sparse lexical matching (TF-IDF) and context-aware reasoning [22, 23]. Vector-based ontology indices are precomputed and stored in ChromaDB, enabling fast retrieval and structured candidate evaluation across ontologies such as MONDO, UBERON, PATO, HANCESTRO, HsapDv, and NCBI Taxonomy.

#### 3.3.2. System Components and Agent Roles

The pipeline is composed of two broad categories of components as shown in **Figure 1**: (i) deterministic pre-processing modules for data acquisition and formatting, and (ii) reasoning agents that perform interpretation, validation, and transformation using LLM-based cognitive workflows.

**Figure 1.**
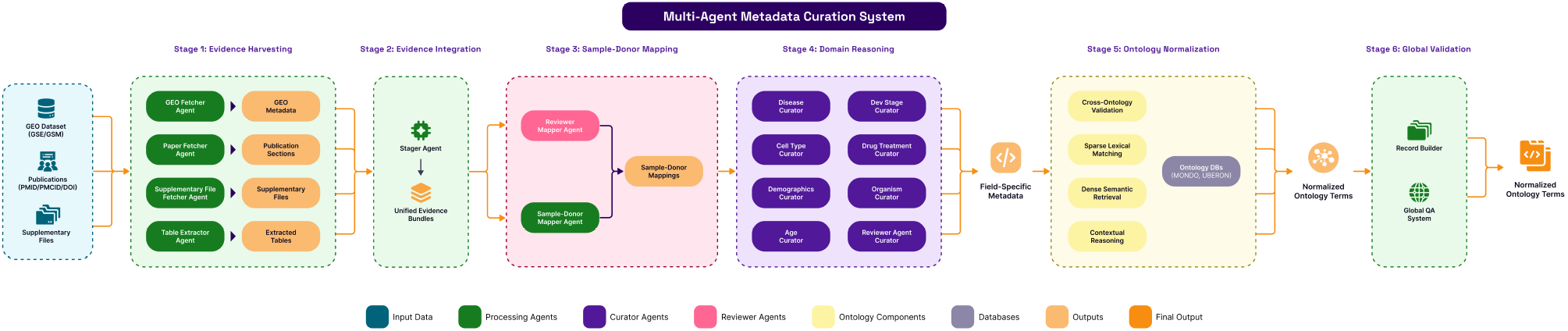
Overview of the Multi-Agent Metadata Curation System: The system ingests GEO datasets, publications, and associated supplementary files using specialized workflows called ‘fetchers’. Curator and reviewer agents generate sample-donor mappings and field-specific metadata, which are normalized to standardized pre-defined ontology terms through semantic and lexical techniques, resulting in normalized, high quality labelled datasets.

##### Deterministic Modules: Data Acquisition and Preparation

These components operate outside the agentic framework. They are designed as fast, reliable scripts or services that run before any AI-based reasoning and prepare the input evidence for downstream consumption:

- **GEO Fetcher:** Retrieves study and sample-level metadata via the NCBI E-utilities API. Extract fields such as title, summary, characteristics, and embedded publication identifiers and validates the data format and structural completeness.
- **Paper Fetcher:** Attempts resolution of associated publications using available identifiers (PMID, PMCID, DOI) via Entrez and Crossref APIs. When full-text access is available, the paper is downloaded, segmented into canonical sections (e.g., Introduction, Methods, Results), and stored as a structured JSON document.
- **Supplementary File Fetcher:** Downloads supplementary data files referenced in the publication, including PDFs, spreadsheets, and text files. Each file is validated for integrity and structure.
- **Table Detection and Extraction Pipeline:** Identifies tables containing pages in PDF supplements using rule-based heuristics (e.g., gridlines, whitespace alignment) and saves them as images. These images are then passed to GPT-4.1 (with vision capability), which extracts tabular data and associated metadata (e.g., page number, table title) as structured JSON. Tables referencing donors or subject-level demographics are explicitly tagged to facilitate downstream mapping.

##### LLM-Based Agents: Cognitive Reasoning and Validation

Once the evidence is assembled, the core reasoning and transformation steps are performed by specialized agents that use GPT-4.1. Each agent implements either chain-of-thought or tree-of-thought workflows, depending on the complexity and ambiguity of the task:

- **Stager Agent:** Merges all available evidence (GEO metadata, publication sections, and parsed tables) into a unified package per sample. Applies consistency checks, flags conflicting information, and preserves source-level attribution.
- **Sample-Donor Mapper Agent:** Uses tree-of-thought reasoning to hypothesize mappings between GSM samples and donor identifiers. Catalogs available mapping evidence, generates candidate hypotheses, evaluates and cross-validates each one, and refines mappings through iterative reasoning steps.
- **Mapper Reviewer Agent:** Independently re-executes the mapping task via an alternate reasoning path. Discrepancies between primary and reviewer output are resolved through multiturn analysis and convergence logic.
- **Curator Agents (8 total):** Each field of interest is assigned to a dedicated curator agent: Disease, Tissue, Cell Type, Organism, Developmental Stage, Age, Demographics, and Drug/Treatment. Each agent interprets field-specific evidence, formulates hypotheses, validates across sources, and performs iterative refinement.
- **Reviewer Agents (Curators):** Mirror each curator agent to independently validate reasoning using an alternative logic path. The final outputs are accepted only upon agreement or reconciled consensus.
- **Ontology Normalization Agent:** For fields that require ontology linkage, this agent performs a multistep process: candidate generation using both dense (embedding-based) and sparse (lexical) retrieval; context-aware scoring that incorporates biological plausibility, definitional alignment, and domain-specific appropriateness; and iterative refinement through reasoning chains that resolve ambiguity and validate consistency with other fields.
- **Record Builder & QA Agent:** Assembles all curated fields into a complete metadata record. Applies integrity checks, verifies biological coherence across fields, resolves internal conflicts, and scores record-level quality.
- **Curation Pipeline**

The system executes metadata curation through a six-stage pipeline, where each stage builds on the outputs of the previous one to progressively transform raw inputs into structured, ontology-normalized sample metadata. Unlike traditional linear processing pipelines, this system incorporates multipath reasoning, iterative validation, and domain-specific heuristics in multiple steps, allowing greater resilience in the face of noisy, ambiguous, or missing data. In the following, we describe the operational flow of each stage in detail.

##### Stage 1: Evidence Harvesting with Validation Trees

Curation begins with automated collection of all available evidence for a GEO series (GSE) and its constituent samples (GSM). This includes:

- **Structured Metadata Retrieval:** GEO metadata is retrieved using the E-utilities API, parsed into per-sample JSON records, and validated for structural integrity.
- **Publication Access Resolution:** The pipeline identifies all available publication identifiers (PMID, PMCID, DOI) and attempts to access through parallel pathways. For each successful route, the content is validated and stored with provenance.
- **Supplementary File Handling:** Supplementary files referenced in publications are downloaded and checked for format, completeness, and relevance.
- **Table Identification & Extraction:** PDF-based supplementary files are parsed to identify tables containing pages using layout heuristics. These pages are rendered as images and sent to a visual LLM (GPT-4.1) for structured extraction. The LLM returns the tabular data along with metadata such as table title, page number, and inferred field types. Tables relevant to the demographics of the subject are explicitly tagged.

##### Stage 2: Evidence Integration with Consistency Reasoning

Once all sources have been harvested, a stager agent constructs evidence bundles per sample. This agent performs a structured inventory of all inputs: GEO metadata, publication sections, and extracted tables and executes the following:

- Checks for intra-source and cross-source consistency,
- Resolves redundancies and source conflicts via rule-based prioritization and contextual cues,
- Flags incomplete or contradictory bundles for further attention.

Each resulting bundle contains all the information available about a single GSM sample, segmented by evidence source, and annotated with metadata confidence levels. This forms the atomic unit of reasoning for the rest of the pipeline.

##### Stage 3: Sample-Donor mapping

A key challenge in GEO metadata is the lack of one-toone mapping between samples (GSM) and biological donors. To address this, a dedicated Sample-Donor Mapper Agent initiates tree-of-thought reasoning over the sample bundle, with the following sequence:

- **Evidence Inventory:** Gathers donor-related fields across GEO, publication body, and tables (e.g., donor IDs, group labels, sex/age pairs).
- **Hypothesis Generation:** Constructs plausible donor identities based on combinations of sample characteristics, donor groupings, and references in tables.
- **Evaluation:** Scores hypotheses using both supporting and contradicting evidence; calculates the confidence per hypothesis path.
- **Cross-Validation:** Validates consistency across evidence types (e.g., ensuring that table-derived donor IDs match GEO-provided labels).
- **Iterative Refinement:** Adjusts mappings based on score differentials, evidence confidence, and overall biological plausibility.

A Reviewer Mapper Agent independently replicates this reasoning using alternate logic, such as preferring tabular information over free text or using stricter consistency criteria. Discrepant outcomes trigger iterative resolution protocols, ensuring donor mappings are only finalized upon consensus.

##### Stage 4: Parallel Domain Reasoning Trees with Independent Validation

Once donor sample associations are established, field-level curator agents activate in parallel to extract domain-specific metadata. Each curator follows a disciplined, multiphase procedure:

- **Evidence Analysis:** Scans all components of the bundle for relevant mentions (e.g., mentions of “bronchoalveolar lavage fluid” for tissue).
- **Hypothesis Formation:** Constructs one or more candidate values based on extracted strings, context cues, and prior domain knowledge.
- **Evaluation:** Applies chain-of-thought reasoning to test biological plausibility, source credibility, and definitional alignment.
- **Cross-Validation:** Confirms whether multiple sources support or contradict the same candidate (e.g., disease mentions in both GEO and publication).
- **Iterative Refinement:** When ambiguity persists, the agent requests clarification subqueries, revisits weak evidence paths, or defers the final decision.

Reviewer agents are run in parallel using different weightage schemes, evidence orderings, or stricter reasoning constraints. Field-level outputs are only accepted when both agents converge or reach a rational reconciliation. All outputs at this stage are expressed in natural language of free text or minimal structured forms (e.g. ‘lung’, ‘left lower lobe’ for tissue), with confidence scores and rationale logs retained.

##### Stage 5: Multi-Candidate Ontology Reasoning with Systematic Exploration

To ensure harmonization between data sets and alignment with external ontologies, curated values undergo normalization through a dedicated ontology reasoning agent. This process includes:

- **Candidate Generation:** Dense Retrieval (Embedding-based semantic search using SapBERT and PubMedBERT embeddings), Sparse matching (TF-IDF-based retrieval capturing lexical and character-level similarities) and Ontology extension (Exploring synonyms, child / parent nodes, and related terms).
- **Contextual Scoring:** Lexical Alignment (Matches between entity mentions and label strings), Semantic Coherence (Conceptual similarity based on definition embeddings) and biological appropriateness (fit within the surrounding metadata context).

The final score combines raw similarity and contextual modulation:

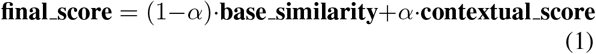

where:

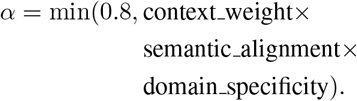

- **Validation & Refinement:** The top-ranked candidates are reviewed through alternate reasoning paths, including boundary tests (e.g., minimal evidence scenarios), crossfield coherence checks, and sensitivity analyzes. Normalized terms include both ontology IDs and human-readable labels.

##### Stage 6: Global Validation and Record Assembly

The final stage composes a unified metadata record per GSM sample. A record builder agent aggregates field-level outputs, validates cross-field logic (e.g., ‘neonatal’ age with ‘adult lung’ tissue), and verifies consistency between curated values and donor-level context. The record is structured into a final output format, including:

- All normalized field values,
- Source-specific evidence attribution,
- Confidence metrics per field,
- Full reasoning provenance.

A QA agent conducts a comprehensive check, including:

- **Completeness Validation:** Are all required fields present?
- **Consistency Validation:** Do field combinations align biologically?
- **Integrity Validation:** Are mappings and identities consistent in the steps?
- **Quality Assessment:** Are confidence scores above the threshold?

This end-to-end pipeline operationalizes a structured, multipath reasoning system over heterogeneous biomedical metadata. Its layered, modular design supports transparency, extensibility, and robustness at every stage from evidence harvesting through ontology-normalized record delivery.

### 3.5. Evaluation Protocol

To systematically assess the quality and robustness of our multi-agent metadata curation system, we designed an evaluation framework grounded in field-specific ground truth comparisons, standard classification metrics, and baseline benchmarking. This section outlines the construction of the evaluation dataset, the metrics used, and the methodology for comparative analysis.

#### 3.5.1 Ground Truth Construction

The evaluation was performed against manually curated metadata records constructed through a dual-curation process. Two independent human annotators, each with biomedical training and curation experience, reviewed each GSM sample and associated evidence to extract structured values for key metadata fields. Disagreements were evaluated by a third expert reviewer through structured evidence-based reconciliation.

Furthermore, we performed a concordance analysis using data from the CellxGene corpus, focusing on key fields such as disease, tissue, tissue type, and gender. All ground truth records in CELLxGENE are normalized to the same ontologies spaces that our system uses (e.g., MONDO for diseases and UBERON for tissues), thus enabling fair comparison.

#### 3.5.2 Evaluation Metrics

We used standard classification metrics to assess the performance of the prediction per field. For each metadata field (e.g. tissue, disease), we calculated:

- **Precision (weighted-macro):** Weighted Fraction of predicted values of the system that corresponded to the ground truth.
- **Recall (weighted-macro):** Weighted Fraction of the ground-truth values correctly recovered by the system.
- **F1-Score (weighted-macro):** Weighted Harmonic mean of precision and recall, averaged across all samples (weighted-macro).
- **Accuracy:** Fraction of exact matches across all samples and fields.

All comparisons were performed per field, per sample, and the metrics were aggregated across the entire evaluation set.

#### 3.5.3. Baseline Configuration

To contextualize the performance of the entire multi-agent system, we implemented a baseline condition using a singlepass GPT-4.1 prompt. This baseline setup was designed to mirror a conventional usage pattern of large language models without agentic structure, retrieval, or iterative reasoning. The configuration was as follows:

- The model received the full evidence bundle (GEO metadata + publication text + tables, flattened) in a single prompt.
- Instructions were given to extract and return metadata fields as structured JSON.
- No chain-of-thought prompting, function calling, or intermediate validation steps were used.
- The raw extracted terms were normalized to their respective ontologies using the same normalization agent used in the agentic approach.

This baseline reflects a simplified zero-shot extraction scenario and serves as a comparison point to highlight the gains achieved through modular reasoning, tree-based exploration, and validation workflows throughout the system.

#### 3.5.4. Evaluation Mode and Protocol

The evaluation was performed using a reproducible script that aligned the predicted records with ground truth using GSM ID, and then evaluated each field independently. Metrics were computed for: each metadata field independently and each sample individually (per-sample weighted macro-scores). All predictions were stored with reasoning logs, provenance traces, and confidence scores to support downstream error analysis. Summary statistics and field-level performance charts are reported in the results section.

## 4. Results

### 4.1. Performance of Agentic System in curating key metadata attributes compared to Baseline (GPT-4.1 Using Single-Pass Prompting)

To evaluate the performance of our multi-agent metadata curation system, we conducted a comparative performance analysis against a simplified single-pass prompting method that used GPT-4.1. Both methods used the same data sources. However, the agentic system (**Figure 1**) integrates multiple specialized agents for data extraction, validation, and ontology normalization, as detailed in the system architecture, while the baseline method uses a single pass extraction.

The evaluation was carried out on a corpus of 78 GEO datasets (**Supplementary Table 2**), consisting of 4652 samples (GSM IDs). In the context of GEO, each GSM ID corresponds to a unique biological sample or experimental unit submitted by the authors with associated high-throughput gene expression data. Expert reviewers manually reviewed the metadata for each sample, validating original label terms, normalized equivalents, and ontology identifiers for both the baseline and agentic workflows. This comprehensive review allowed the computation of precision, recall, F1 score, and accuracy for both systems.

**Figure 2** presents a comparison of performance metrics between the two systems. The agentic system consistently outperformed the GPT-4.1 zero-shot baseline for all evaluated fields, particularly in context-dependent metadata categories.

**Figure 2.**
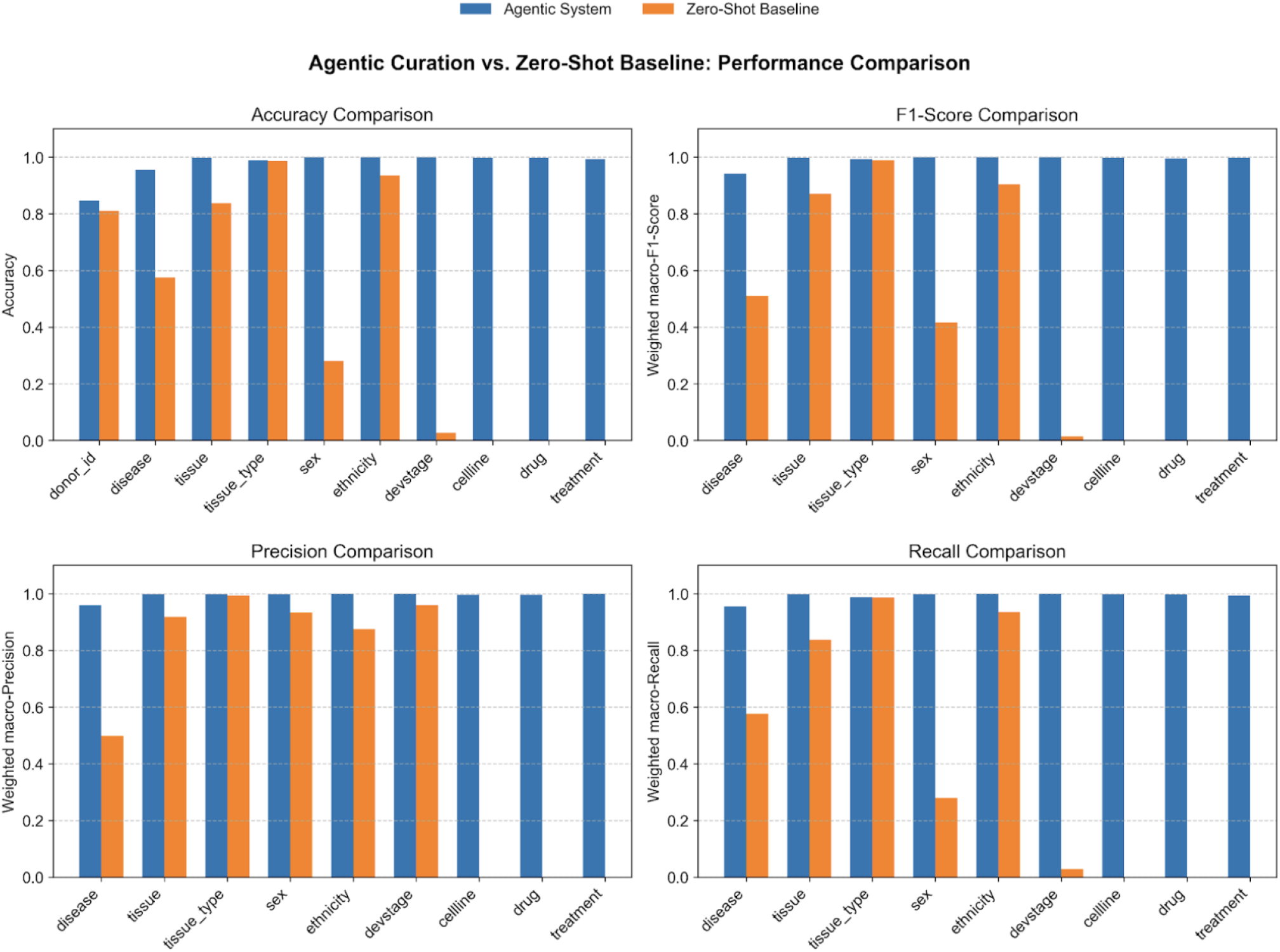
Comparative performance of the agentic metadata curation system versus a simplified GPT-4.1 zero-shot baseline across four metrics - precision, recall, F1-score, and accuracy. The agentic system consistently outperforms the baseline, particularly in context-dependent fields such as disease, tissue, and donor-related attributes, demonstrating its effectiveness in structured biomedical metadata extraction and normalization.

#### Precision

Both systems demonstrated high precision in structured fields such as organism (data not shown) and tissue type. However, the agentic system maintained greater than 0.98 precision in nearly all fields, reflecting its robust validation and ontology resolution steps.

#### Recall

The agentic system achieved a recall greater than

0.95 for disease, tissue, and gender. In contrast, the zeroshot baseline struggled with context-heavy fields such as developmental stage and donor ID, with recall values as low as 0.02 for developmental stage.

#### F1-Score and Accuracy

The overall F1-scores and accuracies mirrored recall trends, affirming the advantage of the agentic system in consistently retrieving complete and correct metadata. Baseline performance dropped significantly for underrepresented or contextually complex labels.

In particular, the donor ID field had the lowest accuracy (0.847) in the agent pipeline. A likely reason for reduced recall for the donor identification field is the presence of corner cases that require further handling, as exemplified by the GSE210272 data set. All samples in GSE210272 have cross references to samples in GSE37147 where the actual donor ids are mentioned. This is a rare corner case that the agentic system is not designed to handle.

Although the agentic system performed better than the zero-shot baseline in most metadata fields, both approaches showed similar results for ethnicity extraction. This similarity occurred because roughly 90% of the samples had ethnicity labels missing in both the original GEO metadata and related publications. Since both systems had limited ground truth data to work with, their comparable performance scores do not accurately represent their true extraction abilities for this attribute.

These results highlight the limitations of a single-pass prompting approach, even with a powerful model like GPT4.1, for comprehensive metadata curation. While the zeroshot baseline can achieve reasonable precision in straightforward cases, it lacks the contextual reasoning and multistep validation required for complex or implicit metadata attributes.

In contrast, the agentic system that leverages specialized agents, and a unified evidence model shows near-human performance in key biomedical metadata fields. This makes it especially suitable for high-stakes applications in biomedical data integration, meta-analysis, and downstream computational pipelines that require harmonized metadata.

### 4.2. Concordance on Labels available on Open Source data (CELLxGENE)

CellxGene provides one of the largest curated repositories of publicly available single-cell data sets, including data from 252 + distinct tissues and 34+ unique assay types. The collection includes samples that span normal, 90 unique diseases in multiple species such as Mus musculus, Macaca mulatta, Callithrix jacchus, and Sus scrofa domesticus. Data for each organism are organized into separate SOMAExperiment objects, which are specialized instances of the broader SOMACollection structure. Each SOMAExperiment contains a data matrix (cells by genes), along with associated cell and gene metadata. The cell level metadata comprises of Identifiers like donor id, metadata like cell type, tissue, sex, disease, development stage, and their corresponding ontology term IDs, sample context like assay, suspension type, demographics like self reported ethnicity, gender and the corresponding ontology term IDs, tissue classification like including tissue type [20]. It must be noted that we added the labels at sample level whereas CZI stores metadata at cell level without any GSE or GSM identifiers. The only way to connect GSM ids to CZI data is by means of donor ids. Additionally, CZI has created unique identifiers for many GEO datasets to make donor IDs unique across all studies in a collection. Therefore, identifying the right DOIs that have donor ids that match with the publication & GSM pages that can be used to measure concordance is a tedious step.

To evaluate the concordance between the extraction of our system’s metadata and the curated labels provided by the Chan Zuckerberg Initiative (CZI) CellxGene platform, we compared four key metadata fields between donor samples matched from 21 unique DOIs, as shown in **Table 1**.

The results summarized in **Table 1** indicate strong agreement between key labels, namely disease, tissue, tissue type, and gender. Specifically, disease and tissue type labels demonstrated perfect concordance (1.0), while tissue labels achieved a high concordance of 0.934, reflecting occasional differences in specificity (e.g., “lung” vs. “bronchiole”). The gender labels showed a concordance of 0.97, suggesting near-complete alignment, with minor discrepancies potentially attributable to missing or ambiguous source annotations.

Additional findings emerged during the concordance evaluation. In the data set associated with DOI 10.7554/eLife.62522, donor IDs D062 (GSM4906338), D122 (GSM4906342), and D231 (GSM4906344) are labeled “normal” in the CZI metadata. However, our system assigned more specific disease terms: respiratory distress syndrome in premature infants for D062 (GSM4906338), anthracosis for D122 (GSM4906342) and inflammatory disease for D231 (GSM4906344). Manual verification of the additional information provided to these donors confirmed that the assignments of the system were accurate, aligned with what would be expected from expert human curation. A trace of how our system annotated the disease term for one of the above mentioned samples is shown in **Figure 3**.

**Figure 3.**
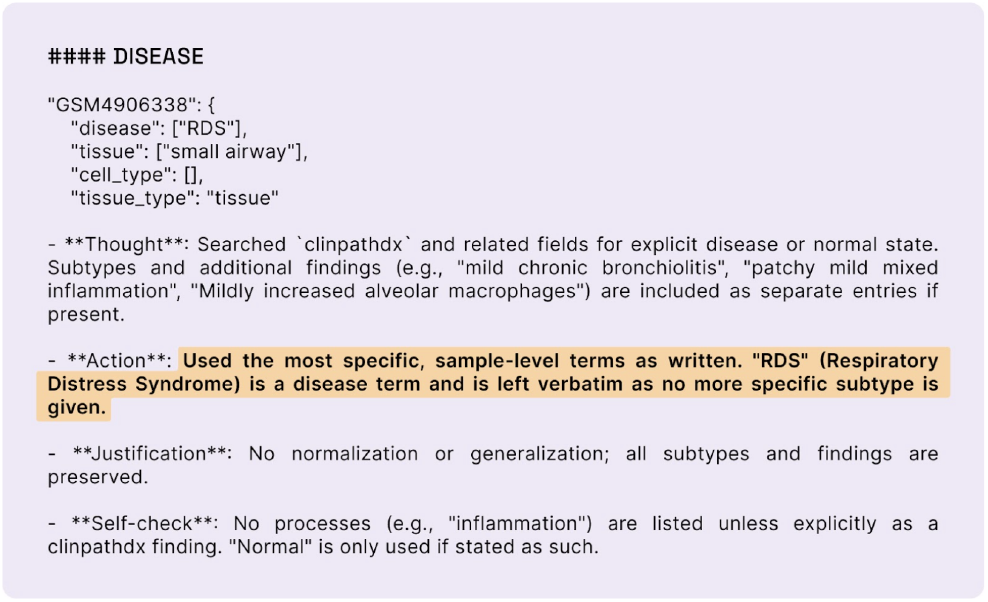
Curation Decision Log for Disease Metadata Extraction from GSM4906338. This panel illustrates the agentic system’s decision-making trace for assigning a disease label. This provenance layer facilitates auditability and high-confidence metadata extraction.

In the same dataset, the tissue term for all donors is listed as “lung” in the CZI metadata. However, our system correctly identified the more specific term “Bronchiole,” as confirmed by manual expert verification. A trace of the call out is shown in **Figure 4**.

**Figure 4.**
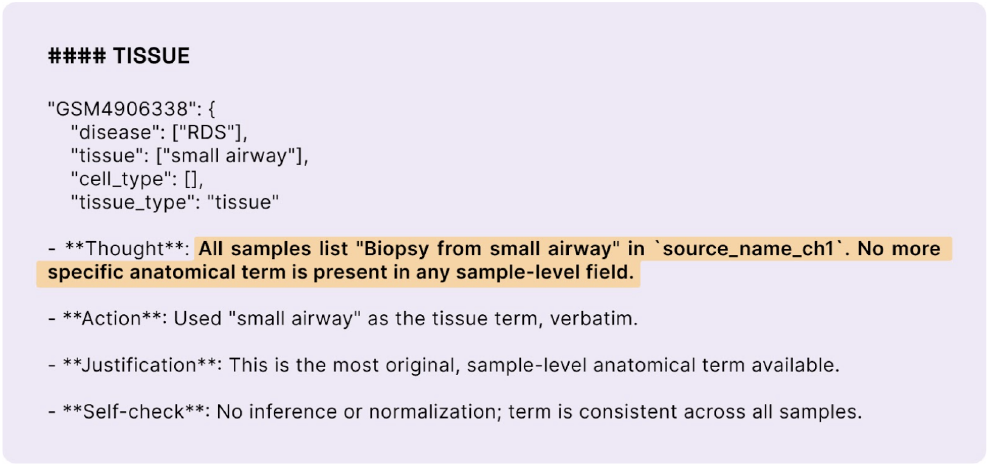
Curation Trace for Tissue Annotation from GEO Sample GSM4906338. This panel documents the agentic curation system’s tissue annotation decision based on sample-level metadata. The system extracted “small airway” from the tissue field, where all samples consistently reported “Biopsy from small airway.” No more granular anatomical descriptor was present. Accordingly, “small airway” was retained verbatim as the most specific tissue term available. No inference, normalization, or substitution was performed.

Similarly, in the data set associated with DOI 10.1016/j.isci.2023.108572, the gender label for the donor ID CV-012 is recorded as “unknown.” In contrast, our system accurately identified and assigned the label “female” to this donor, as confirmed by manual expert review. These instances demonstrate our system’s ability to extract context-aware metadata with expert-level precision, especially in cases where source annotations are overly broad or incomplete. A trace of the call out made by our agentic system is shown in **Figure 5**.

**Figure 5.**
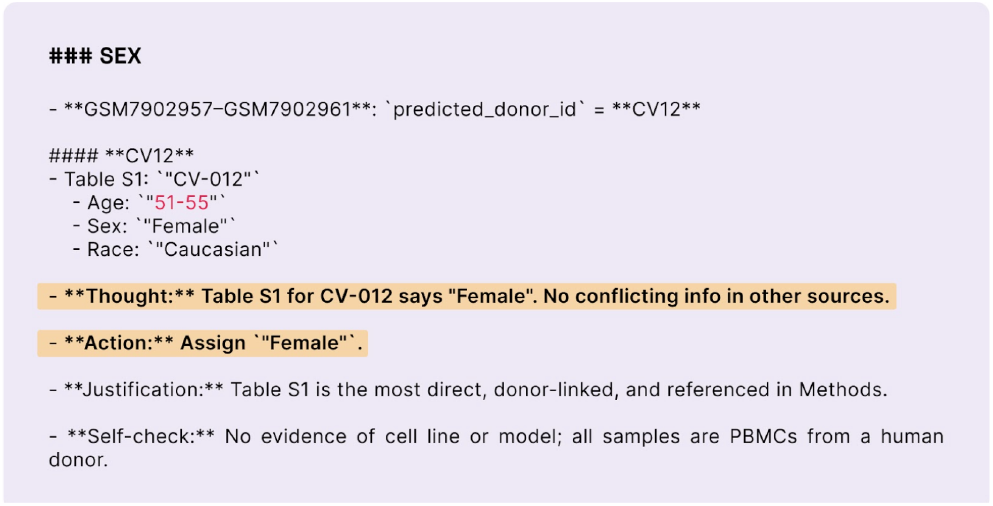
Curation Trace for Sex Annotation of Donor CV12. This panel details the annotation logic for assigning biological sex to samples GSM7902957 - GSM7902961, all linked to donor CV12 via predicted donor id. Table S1 identifies donor “CV-012” as female, with no conflicting sex data from other sources. Therefore, the agentic system assigned “Female” as the sex value. The justification relies on Table S1 as the most direct and study-authored reference, explicitly tied to donor metadata.

## 5. Discussion

Public repositories such as ArrayExpress, Single Cell Portal (SCP), PRIDE, GEO, etc. serve as critical resources for the dissemination of genomic and transcriptomic data in biomed- ical research. However, the integration of datasets across studies is impeded by the lack of standardized, machinereadable metadata, particularly for key biological descriptors such as disease, tissue, sampling site, experimental conditions, etc. Although many GEO entries are linked to peerreviewed publications, these do not resolve the fundamental issue of metadata interoperability for computational reuse. To address these limitations, various strategies have been proposed, including expert-driven manual curation, automated approaches that take advantage of natural language processing (NLP), and inference of metadata from expression profiles for fields of interest such as gender. Manual curation remains the most reliable method in terms of quality assurance, and tools such as GEOMetaCuration have been developed to facilitate this process through a structured workflow involving multi-curator annotation and consensus generation [24].

Although manual curation is often considered the most reliable approach to ensuring high-quality metadata, it presents significant challenges in terms of scalability and time efficiency. This method is generally effective when the number of metadata fields to be extracted and standardized using predefined ontologies is typically limited to 20 or fewer attributes. The curation workflow typically involves assigning the same task (e.g., annotating 20 sample-level fields) to two independent curators working in a blinded fashion, followed by a reconciliation phase in which discrepancies are resolved through discussion. Although this process helps ensure accuracy and consistency, it substantially increases turnaround time and is poorly suited for applications such as machine learning, where large volumes of curated data are needed, often involving more than hundreds of sample-level fields.

Our internal estimates indicate that curating approximately 22,000 samples from GEO (for 12 metadata attributes for each sample) and linked to 211 publications requires around four weeks of effort by a three-member team. Despite this significant investment of time and resources, the resulting dataset only marginally meets the level of granularity required for machine learning applications. Furthermore, any increase in the number of metadata attributes per sample leads to a proportional increase in turnaround time, underscoring the scalability limitations of manual curation in highdimensional data contexts.

To overcome the limitation of manual curation, we developed a multiagent artificial intelligence system to automate metadata curation for Gene Expression Omnibus (GEO) datasets at scale. The system employs a six-stage pipeline architecture that implements tree-of-thought and chain-of-thought reasoning frameworks to extract, validate, and standardize biological metadata from heterogeneous sources while maintaining quality control through multipath validation.

This computational approach uses specialized AI agents with cognitive reasoning architectures for metadata extraction and curation. The modular design enables parallel processing with sequential quality checkpoints. The architecture incorporates reasoning trees where each decision point branches into multiple pathways with independent evaluation and validation, representing a shift from linear processing to multi-path exploration.

The system leverages GPT-4.1 for reasoning-intensive tasks. Ontology normalization uses specialized biomedical embeddings (fine-tuned SapBERT and PubMedBERT) for dense semantic retrieval and sparse representations of TF-IDF for lexical matching. Target ontologies include MONDO (diseases), UBERON (anatomical structures), Cell Ontology (cell types), PATO (phenotypic qualities), HANCESTRO (human ancestry), Human Developmental Stages Ontology (HsapDv) and NCBI Taxonomy (organisms). Vector indices use ChromaDB for ontology term retrieval with precomputed embedding storage.

The evaluation of the agentic system for extracting and normalizing metadata across 78 GEO datasets demonstrates promising performance, particularly in its ability to process more than 10 distinct metadata fields with high accuracy. The overall recall rates indicate that the system is effective in extracting original label terms, generating normalized terms, and linking them to appropriate ontology identifiers across a diverse set of biological and clinical attributes.

Perfect recall (1.0) was achieved for organism-related fields, including original labels, normalized terms, and ontology mappings, underscoring the robustness of the system in handling well-defined, taxonomically stable metadata. Similarly, demographic attributes such as sex and ethnicity, as well as treatment-related fields, exhibited near-perfect recall (greater than 0.99), indicating that these categories are consistently represented in the source data and are well supported by the normalization logic and reference ontologies used.

The system also performed strongly in the tissue and disease-related fields, achieving recall rates greater than 0.95 for original labels, normalized terms, and ontology identifiers. This shows its ability to accurately handle both standardized biomedical terminology and diverse user-provided annotations with minimal information loss.

From a scalability perspective, the agentic approach enables rapid processing of thousands of GEO datasets and can be readily extended to internal data repositories or other sources such as SCP or ArrayExpress with minimal adaptation. Unlike manual efforts that require repeated domain-specific review of each label, the agentic system amortizes this cost through reusable logic, curated reference lists, and intermediate self-checks that support traceability and auditability. Furthermore, its modular design allows for easy integration into ETL (Extract, Transform, Load) pipelines and LLM-based downstream tools, reducing the dependency on bespoke, domain-expert-driven annotation cycles.

We adopted a publicly available metadata schema developed by the Chan Zuckerberg Initiative (CZI), version 5.2.0, originally designed for single-cell datasets, and extended it to incorporate additional attributes such as data type, cell line, drug, and treatment. These specific metadata fields were chosen due to their prevalence across a substantial proportion of datasets hosted in major public repositories, including ArrayExpress, GEO, etc. The modified schema enables a more comprehensive representation of experimental conditions commonly reported in datasets present on ArrayExpress, GEO, etc. In addition, we identified several opportunities to improve the existing schema by incorporating new fields, including age, age units, drug dosage, and sampling site. We propose a revised schema that explicitly separates tissue terms from sampling site descriptors. This separation supports distinct downstream applications: the broad tissue term enhances dataset findability through standardized ontology mappings, while the sampling site provides critical contextual information relevant to the biological interpretation of the sample. Our evaluation indicates that these additional fields also achieve a very high level of recall, as confirmed by expert review (data not shown).

Overall, our architected solution provides a scalable and robust system that leverages state-of-the-art LLMs for metadata harmonization and demonstrates high reliability across a wide range of metadata fields. It supports downstream applications such as meta-analysis and machine learning by enabling high-throughput metadata curation with expert-level accuracies. Future improvements of the system will focus on resolving rare edge cases, enhancing inference across linked datasets, and expanding support for underrepresented metadata types through integration with external knowledge sources, particularly for identifiers not explicitly encoded within individual datasets.

## Acknowledgments

The authors thank Gaurang Mahajan, Rahul Tyagi, Shubhra Agrawal, and Swetabh Pathak for their valuable feedback and support.

